# An RNA-binding tropomyosin recruits kinesin-1 dynamically to *oskar* mRNPs

**DOI:** 10.1101/067876

**Authors:** Imre Gáspár, Vasily Sysoev, Artem Komissarov, Anne Ephrussi

## Abstract

Localization and local translation of *oskar* mRNA at the posterior pole of the *Drosophila* oocyte directs abdominal patterning and germline formation in the embryo. The process requires recruitment and precise regulation of motor proteins to form transport-competent mRNPs. We show that the posterior-targeting kinesin-1 is loaded upon nuclear export of *oskar* mRNPs, prior to their dynein-dependent transport from the nurse cells into the oocyte. We demonstrate that kinesin-1 recruitment requires the *Dm*Tropomyosin1-I/C isoform, an atypical RNA-binding tropomyosin that binds directly to dimerizing *oskar* 3’UTRs. Finally, we show that a small but dynamically changing subset of *oskar* mRNPs gets loaded with inactive kinesin-1 and that the motor is activated during mid-oogenesis by the functionalized spliced *oskar* RNA localization element. This inefficient, dynamic recruitment of Khc decoupled from cargo-dependent motor activation constitutes an optmized, coordinated mechanism of mRNP transport, by minimizing interference with other cargo-transport processes and between the cargo associated dynein and kinesin-1.

## Introduction

Within cells, organelles, diverse macromolecules and complexes depend on a small set of cytoskeleton-associated motor proteins to achieve their proper distributions Targeted delivery is ensured by cargo-associated guidance cues that are responsible for recruiting the appropriate mechanoenzyme (Hirokawa, Niwa et al., 2010). Such actively transported cargoes include mRNAs, whose asymmetric localization and local translation within cells are essential for various cellular functions, such as migration, maintenance of polarity and cell fate specification (Medioni, Mowry et al., 2012). In the case of messenger ribonucleoprotein (mRNP) particles, the guidance cues are the mRNA localization elements (LE) that suffice to drive localization of the RNA molecule that contains them (Marchand, Gaspar et al., 2012). A few LEs, their RNA binding proteins (RBP) and the factors that link them to the mechanoenzyme have been well characterized (Bullock, Ringel et al., 2010, Dienstbier, Boehl et al., 2009, Dix, Soundararajan et al., 2013, Niedner, Edelmann et al., 2014). In these cases, the entire localization process is driven by a single type of motor. Other mRNAs, such as *Xenopus laevis Vg1* (Gagnon, Kreiling et al., 2013) and *Drosophila melanogaster oskar (oskar)* (Clark, Meignin et al., 2007, Jambor, Mueller et al., 2014, Zimyanin, Belaya et al., 2008) rely on the coordinated action of multiple motor proteins - cytoplasmic dynein and kinesin-1 and -2 family members - for their localization within developing oocytes.

*oskar* mRNA encodes the posterior determinant Oskar protein, which induces abdomen and germline formation in a dosage-dependent manner in the fly embryo (Ephrussi & Lehmann, 1992). *oskar* mRNA is transcribed in the nurse cells of the germline syncytium and transported into the oocyte, similarly to e.g. *bicoid* and *gurken* mRNAs (Ephrussi, Dickinson et al., 1991, Kim-Ha, Smith et al., 1991). This first step of *oskar* transport is guided by a well described LE, the oocyte entry signal found in the 3’UTR of the mRNA (Jambor et al., 2014), which is thought to recruit the Egl-BicD-dynein transport machinery (Clark et al., 2007, Jambor et al., 2014). In the oocyte, *oskar* mRNA localization to the posterior pole is mediated by kinesin-1 (Brendza, Serbus et al., 2000, Loiseau, Davies et al., 2010, Zimyanin et al., 2008). This second step of *oskar* transport requires RNA splicing (Hachet & Ephrussi, 2004), which assembles the spliced localization element (SOLE) and deposits the exon junction complex (EJC) on the mRNA, to form a functional EJC/SOLE posterior targeting unit (Ghosh, Marchand et al., 2012). The EJC/SOLE was shown to be crucial for maintaining efficient kinesin-1 dependent transport of *oskar* mRNPs within the oocyte(Ghosh et al., 2012, Zimyanin et al., 2008), which is essential for proper localization of the mRNA to the posterior pole.

In a forward genetic screen, a group of *DmTm1* (formerly *DmTmII*) mutants (*Tm1*^*gs*^) was identified in which *oskar* mRNA accumulation at the posterior pole of the oocyte fails (Erdelyi, Michon et al., 1995, Zimyanin et al., 2008) (Fig 1A and B). Although the small amount of Oskar protein produced at the posterior pole is sufficient for embryo progeny of *Tm1*_*gs*_ homozygous females to form an abdomen and develop into adult flies, it is insufficient to induce primordial germ cell formation. Consequently, the *Tm1*_*gs*_ progeny are sterile, resulting in a so-called ‘grandchildless’ phenotype (Erdelyi et al., 1995). It was subsequently demonstrated that the microtubule mediated intra-ooplasmic motility of *oskar* mRNPs is affected in *Tm1*_*gs*_ mutants (Zimyanin et al., 2008) (Table S1), similar to what has been observed when kinesin-1 is absent (Zimyanin et al., 2008). Although there is biochemical evidence that kinesin-1 associates with *oskar* mRNPs (Sanghavi, Laxani et al., 2013), what mediates the association of the motor with the mRNA, and where in the egg-chamber this occurs is not known.

**Figure 1:**
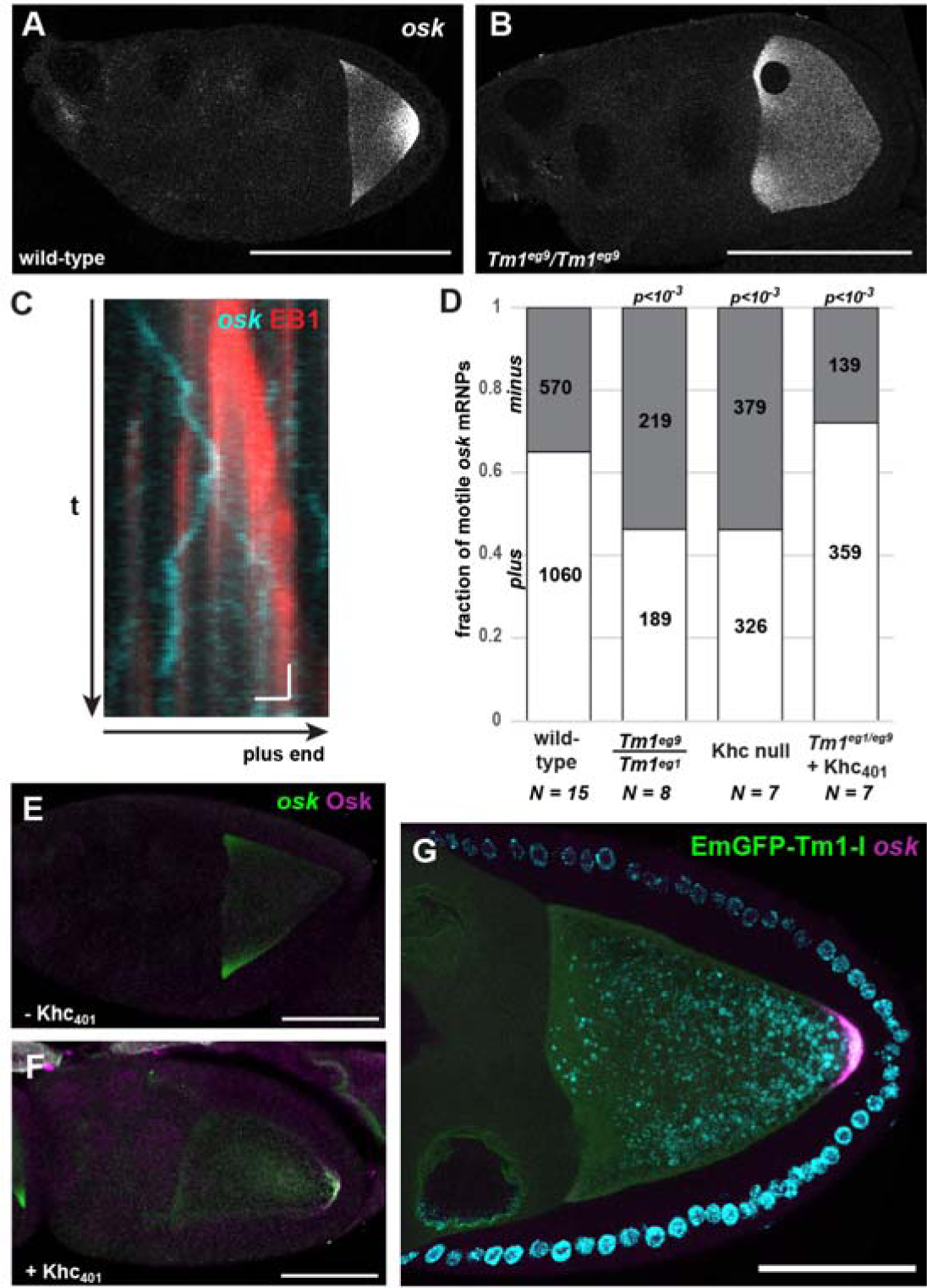
Effect of *Tm1*^*gs*^ on *oskar* transport. (A, B) Localization of *oskar* mRNA in wild-type (A) and in *Tm1*^*eg9*^/*Tm1*^*eg9*^ (B) egg-chambers. Kymograph of *oskMS2*-GFP mRNPs (cyan) travelling along polarity marked MTs (red, EB1 protein) in an *ex vivo* ooplasmic preparation. Scale bars represent 1 s and 1 µm, respectively. (D) Distribution of *oskMS2*-GFP mRNP runs towards plus (white) and minus ends (grey). Numbers within the bars indicate number of runs. P value of Chi^2^ test against wild-type is indicated above each bar. (E, F) *oskar* mRNA (green) and Oskar protein (magenta) distribution in *Tm1*^*eg1*^/*Tm1*^*eg9*^ egg-chambers expressing *oskarMS2(6x)* (E) or *oskarMS2(6x)* and Khc_401_-MCP (F). (G) *oskar* mRNA (magenta) distribution in *Tm1*^*eg1*^/*Tm1*^*eg9*^ egg-chambers rescued with EmGFP-Tm1-I (green). Scale bars represent 50 µm. See also Figs S1 and S4 and Movie S1.

Here, we demonstrate that DmTm1-I/C, a product of the *DmTm1* locus, is an RNA binding tropomyosin that recruits kinesin heavy chain (Khc) to *oskar* mRNA molecules upon their nuclear export in the nurse cells. We show that Khc recruitment to *oskar* RNPs is transient and dynamic, and that this dynamic recruitment depends on the presence of DmTm1-I/C. Our data indicate that, during mid-oogenesis, the EJC/SOLE triggers kinesin-1 activity, which drives localisation of *oskar* mRNA to the posterior pole of the oocyte.

## Results

### Tm1-I maintains kinesin-1 on *oskar* mRNA

To obtain mechanistic information regarding the motility defect in *Tm1*_*gs*_ oocytes, we developed an *ex vivo* assay that allows co-visualization of MS2-tagged *oskar* mRNPs and polarity-marked microtubules (MTs) in ooplasm and determination of the directionality of *oskar* mRNP runs (Fig 1C, Fig S1A and Movie S1), thus giving insight into the identity of the motor(s) affected by the *Tm1*_*gs*_ mutations. Using this assay, we found that in wild-type ooplasm plus end-directed runs of *oskMS2* RNPs dominated about two to one over minus end-directed runs (Fig 1D). Plus-end dominance was lost both in ooplasm lacking Khc and in extracts prepared from *Tm1*_*gs*_ mutant oocytes (Fig 1D). This indicates that plus end-directed, Khc-mediated motility is selectively compromised in the *Tm1*^*gs*^ mutants. The remaining plus end directed runs might be due to residual kinesin-1 activity, to other plus end-directed kinesins, or to cytoplasmic dynein, which has been shown to mediate the bidirectional random walks of mRNPs along MTs (Soundararajan & Bullock, 2014). To test if the loss of Khc activity might be the cause of *oskar* mislocalization in *Tm1*^*gs*^ oocytes, we tethered a minimal Khc motor, Khc_401_ (Sung, Telley et al., 2008, Telley, Bieling et al., 2009), to the MS2-tagged *oskar* mRNPs. Co-expression of Khc_401_–MCP and *oskMS2* restored the plus end dominance of *oskar* mRNP runs (Fig 1D), as well as localization of *oskar* mRNA (Fig 1E and F), indicating that Tm1 acts upstream of Khc and that loss of kinesin-1 activity might cause *oskar* mislocalization in *Tm1*^*gs*^ mutants.

The *DmTm1* locus encodes at least 17 different transcripts and 16 different polypeptides (Fig S2A). By performing semi-quantitative RT-PCR analysis, we found that the transcripts of Tm1-C, I and H are selectively missing or their amount is greatly reduced in *Tm1*^*eg1*^ and *Tm1*^*eg9*^ homozygous ovaries, respectively (Fig S2B and C). Furthermore, an EmGFP-Tm1-I transgene expressed in the female germline rescued *oskar* mislocalization (Fig 1G) and the consequent grandchildless phenotype of *Tm1*^*gs*^ mutants (all female progeny (n>20) contained at least one ovary with developing egg-chambers). This indicates that the Tm1-I/C isoform is essential for *oskar* mRNA localization.

To determine whether the reduction in Khc-dependent *oskar* mRNP motility in *Tm1*^*gs*^ oocytes is due to insufficient kinesin-1 recruitment or to insufficient motor activity we analysed the composition of *oskar* mRNPs in ooplasms of flies co-expressing either Khc-EGFP or EmGFP-Tm1-I and *oskMS2*-mCherry, *ex vivo* (Movie S2 and S3). We first performed an object-based colocalization analysis of single snapshot images corrected for random colocalization (Fig S1E-H). This revealed that both Khc-EGFP (Fig 2A and Movie S2) and EmGFP-Tm1-I (Fig 2B Movie S3) are recruited to a small but significant fraction of *oskMS2*-mCherry mRNPs (1-4%, Fig 2C), indicating that the two proteins are components of *oskar* transport particles (Fig 2C). Furthermore, the association of Khc-EGFP with *oskMS2*-mCherry mRNPs was significantly reduced (2-4 fold) in *Tm1*^*gs*^ mutant ooplasms (Fig 2D and Fig S1H), indicating that the observed motility defects in *Tm1*^*gs*^ mutant oocytes are due to insufficient kinesin-1 recruitment to *oskar* mRNPs.

**Figure 2:**
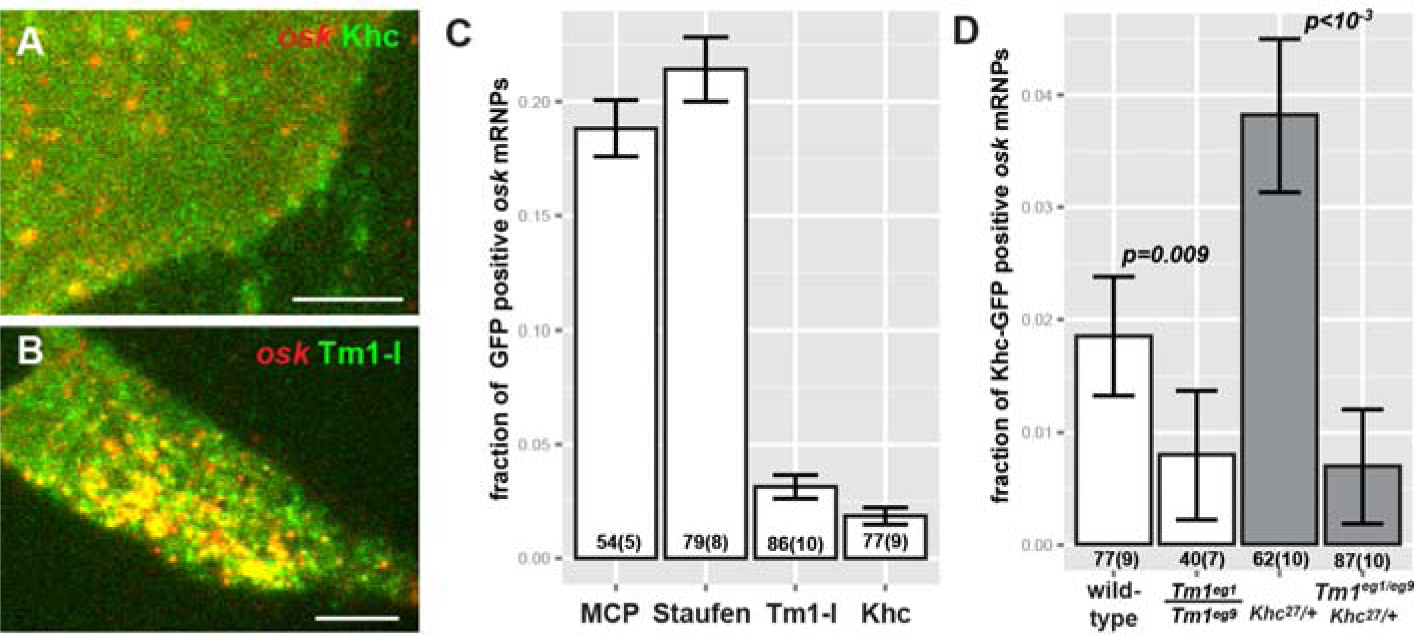
Composition of *oskar* mRNPs ex vivo. (A, B) Colocalization of *oskMS2*-mCherry with Khc-EGFP (A) or with EmGFP-Tm1-I (B) in *ex vivo* ooplasmic preparations. (C) Fraction of *oskMS2*-mCherry mRNPs located non-randomly within a 200 nm distance of one of the indicated GFP tagged protein particles in *ex vivo* ooplasmic preparations. All values are significantly different from zero (p<10^−3^, one sample t-test). (D) Fraction of *oskMS2*-mCherry mRNPs co-localizing (max. 200 nm) non-randomly with Khc-EGFP particles in wild-type and in *Tm1*^*eg1*^/*Tm1*^*eg9*^ ooplasms in presence of two (white) or one (grey) copy of endogenous Khc. (C, D) P values of two sample t-tests are indicated above the relevant bar pairs. Numbers indicate the number of particle clusters (160 mRNPs in each) and the number of preparations (in brackets) analysed. Error bars represent 95% confidence intervals. See also Fig S1 and Movies S2 and S3.

In a complementary approach, we analysed Khc and Tm1-I colocalization with *oskMS2-* EGFP in consecutive images of entire time series. This assay estimates the probability of random co-localization in a different manner (Fig S3A). For the analysis, we made use of flies expressing endogenously tagged Khc and Tm1-I/C, in which virtually all molecules of interest are labelled (Fig 3A and B). This revealed that in *Khc*^*mKate2*^ homozygous ooplasmic extracts nearly 50% of motile *oskMS2-*EGFP mRNPs are associated with Khc during at least half of the recorded trajectories (Fig 3C), close to the proportion of plus-end directed runs (65%) (Fig 1D). In contrast, only ~15% of non-motile mRNPs are associated with Khc during their trajectories. The fact that at any given moment most *oskar* particles are stationary (Movie S1, Table S1)(Gaspar, Yu et al., 2014, Ghosh et al., 2012, Zimyanin et al., 2008) indicates that, in accordance with the analysis of snapshot images (Fig 2D), the majority of *oskar* mRNPs are not in complex with Khc. Our examination of Khc association with RNPs in *Tm1*^*gs*^ mutant extracts revealed that it is equally low in the motile and non-motile mRNP populations, and that it is considerably below that observed in the wild-type control (Fig 3C). This confirms our analysis of snapshot images and demonstrates that Tm1-I/C is required for efficient loading of Khc on *oskar* mRNPs. Furthermore, our finding that approximately 20% of *oskar* mRNPs are stably associated with mCherry-Tm1-I/C irrespective of their motility (Fig 3B and D) indicates that in the oocyte, like Staufen (Fig 3E), Tm1-I/C is a component of *oskar* transport particles.

**Figure 3:**
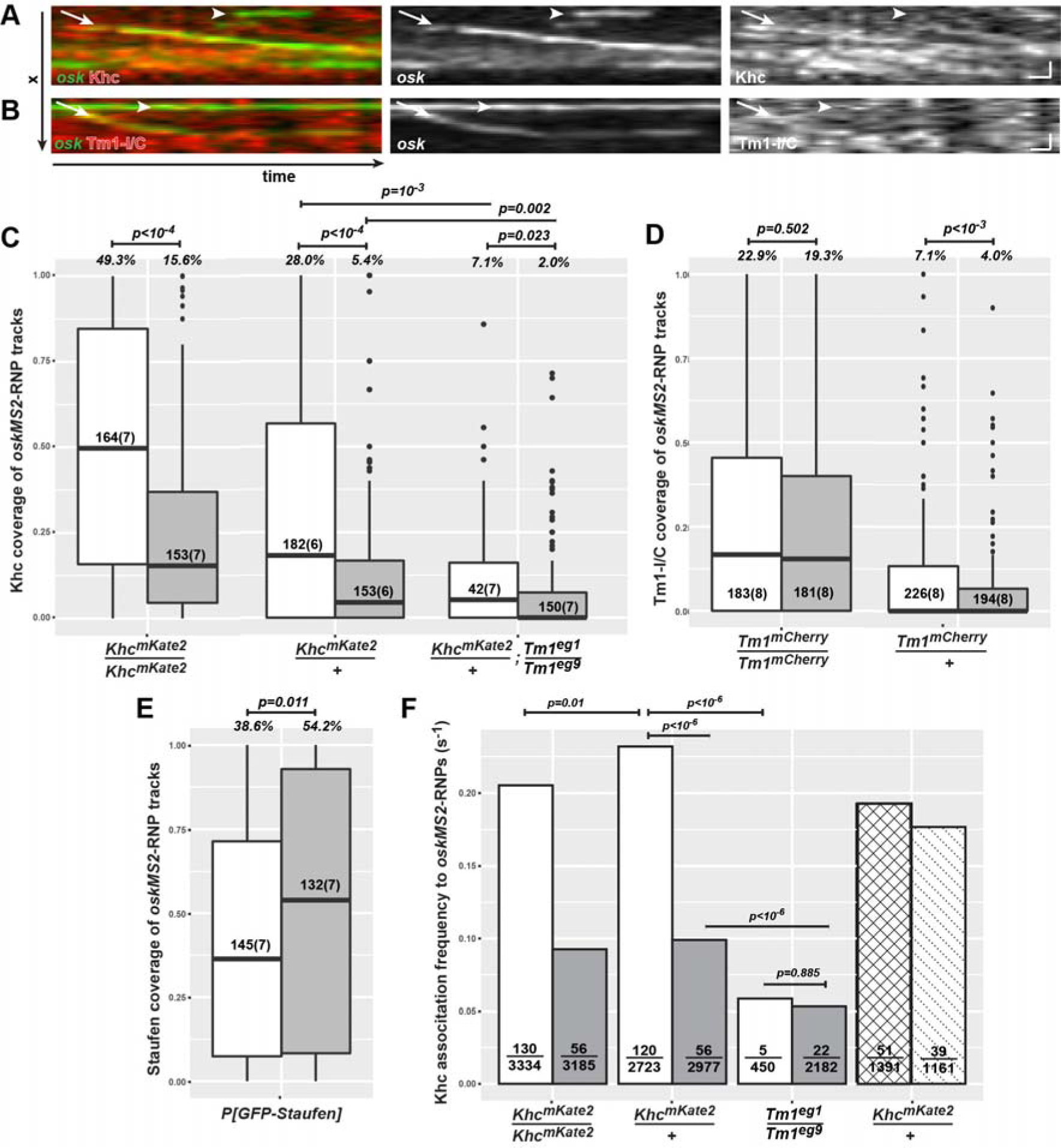
Dynamic composition of *oskar* mRNPs *ex vivo*. (A, B) Kymographs of *oskMS2*-GFP mRNPs (green) associated with Khc-mKate2 (A, red) and mCherry-Tm1-I/C (B, red) *ex vivo*. Arrows indicate motile RNPs in stable complex with Khc (A) or Tm1-I/C (B), the arrowheads point to non-motile *oskMS2*-RNPs showing no obvious accumulation of the tagged protein. Scale bars represent 1 µm and 1 second, respectively. (C-E) Relative Khc-mKate2 (C), mCherry-Tm1-I/C (D) and GFP-Staufen (E) coverage of motile (white) and non-motile (grey) *oskMS2*-GFP trajectories. Numbers within the boxes indicate the number of trajectories and the number of ooplasms (in brackets) analysed. Percentages above the plots show the fraction of RNPs that were found stably and reliably associating with the indicated protein (for at least half of the duration of the trajectory, p<0.01, binominal distribution, see also Fig S3C). P values of pairwise Mann-Whitney U tests are indicated above the boxplots. (F) Frequency of Khc-mKate2 appearance on motile (white), before (motility-primed, checked) and after (dotted) the onset of motility, and non-motile (grey) *oskMS2*-GFP trajectories (see also Fig S3D). Fractions within the bars indicate number of association events that lasted longer than a single frame over the total number of frames analysed. Indicated P values show results of pairwise Fisher’s exact test. The Khc association frequency observed on motility-primed (checked) and moving (dotted) RNPs is not significantly different from wild-type motile RNP controls (p>0.01). See also Fig S3

## Kinesin-1 associates dynamically with *oskar* RNPs

During stage 9 of oogenesis, half of all *oskar* mRNA molecules in the oocyte translocate to the posterior pole (Gaspar et al., 2014). Since only 15% of *oskar* RNPs are in complex with Khc at any given moment (Fig 3C), this implies that kinesin-1 must dynamically redistribute within the RNP population. Our further analysis of time-lapse series, quantifying the degree of Khc association with *oskar* RNPs prior to the onset of motility, revealed a slight increase in Khc occupancy on *oskar* mRNPs roughly 4-5 seconds before the start of runs (Fig S3B). We also measured the frequency of the kinesin-1 and *oskar* association events and observed that motile or motility-primed mRNPs associate with a Khc signal about once every 5 seconds (~0.2 s^−1^, Fig 3F) in wild-type ooplasm. This frequency decreased to ~0.1 s^−1^ in the case of wild-type, non-motile RNPs and dropped to ~0.05 s^−1^ when Tm1-I/C was absent (Fig 3F). This observation, the low degree of Khc association we observe in *Tm1*^*gs*^ mutant ooplasm and the interaction of Tm1-I/C with *oskar* mRNPs (Figs 2C and 3D) indicates that Tm1-I/C acts in recruiting the kinesin motor to the mRNA. The failure of this recruitment explains the greatly reduced number of long, unidirectional – in particular the plus-end directed - runs of *oskar* mRNPs in the absence of Tm1-I/C (Fig 1D, Table S1)(Zimyanin et al., 2008).

### Tm1-I/C is in complex with Khc and *oskar* mRNA and directly binds the *oskar* 3’UTR

If Tm1-I/C is indeed responsible for Khc recruitment to *oskar* mRNPs, these molecules should be in complex with one another. To test this hypothesis, we performed immunoprecipitations from ovarian lysates. Similar to what has been reported previously *in vitro* (Veeranan-Karmegam, Boggupalli et al., 2016), we detected that Khc specifically co-immunoprecipitated with the EmGFP-Tm1-I bait *in vivo* (Fig 4A). Also, we found Staufen, but none of the other tested *oskar* RNP components (Bruno, Y14, BicD or dynein) in the eluate (Fig 4A). Although such an *en mass* co-immunoprecipitation analysis lacks spatiotemporal resolution, it shows that Tm1-I/C, kinesin-1 and Staufen indeed form complexes that are very likely maintained by – not necessarily direct-protein-protein interactions.

**Figure 4:**
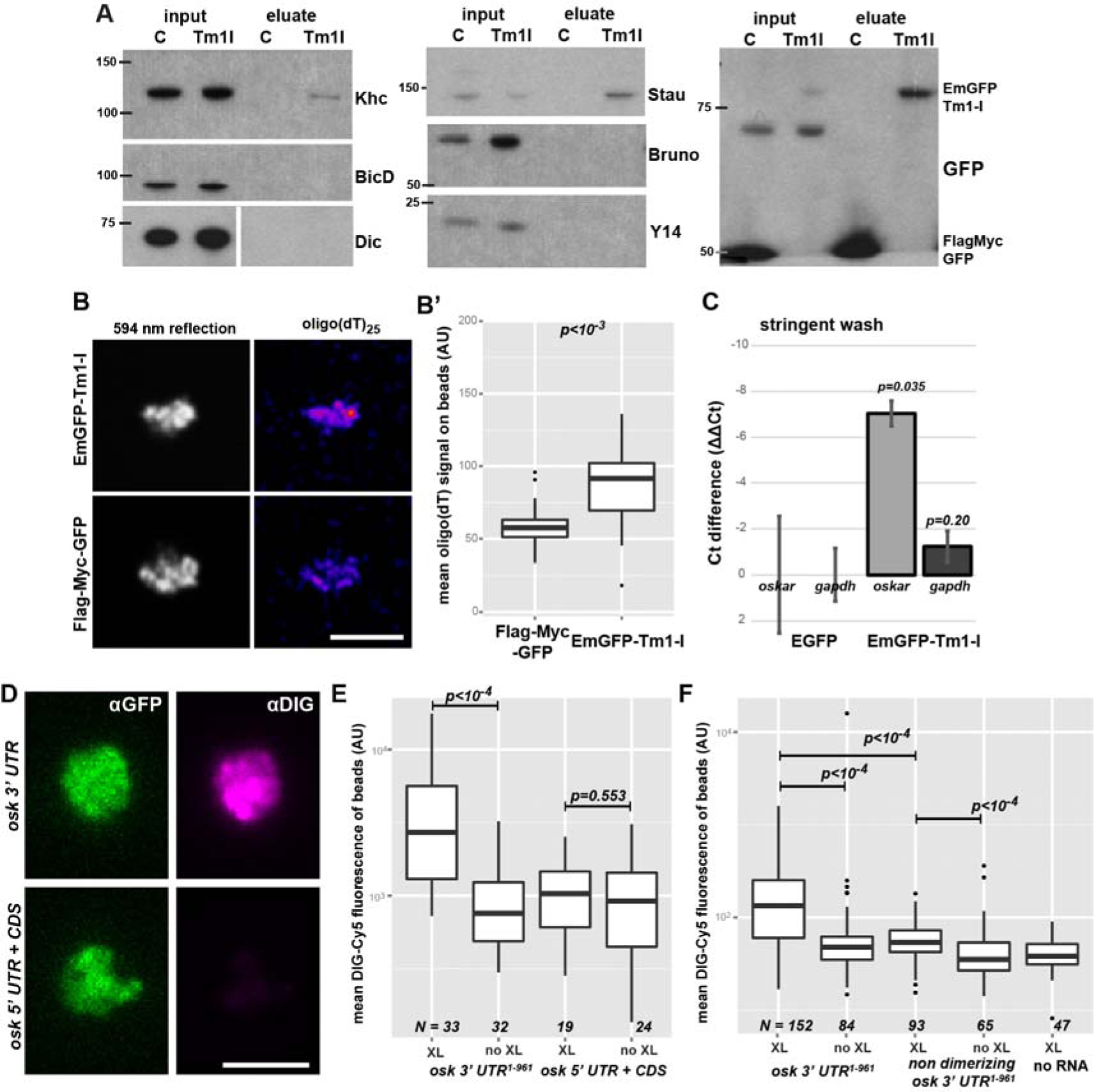
Tm1-I/C binds RNA and Khc. (A) Western blots of *oskar* mRNP components (Staufen, Bruno and Y14) and motor associated proteins (Khc, BicD, Dic) co-immunoprecipitated with EmGFP-Tm1-I from ovarian lysates. Protein marker bands and their molecular weight in kDa are indicated. (B) Signal developed with oligo(dT)^25^-TexasRed probes on GFP-Trap_M beads binding EmGFP-Tm1-I or FlagMycGFP after denaturing washes. *594 nm reflection* marks bead boundaries. (B’) Quantification of mean oligo(dT) signal measured on beads. (c) qRT-PCR of EmGFP-Tm1-I and control Flag-Myc-GFP and bead alone (*w*^*1118*^) bound *oskar* and *gapdh* RNAs after stringent washes (mean ± s.e.m., N = 3). (D) Images of beads binding to EmGFP-Tm1-I (green) and DIG labelled *in vitro* transcribed RNA fragments (magenta). (E, F) Mean DIG-Cy5 fluorescence measured on beads capturing the RNA fragment, with or without UV cross-linking, indicated below the charts. P values of pairwise Mann-Whitney U tests are indicated. In panel F, none of the non-cross-linked samples differ significantly from the no RNA control (p>0.05). Numbers below the plots indicate the number of beads analysed.

In a screen to identify proteins bound directly to mRNAs in early *Drosophila* embryos, we isolated an isoform-non-specific Tm1 peptide (Sysoev, Fischer et al., 2016). By immunoprecipitating EmGFP-Tm1-I from lysates of embryos exposed to 254nm UV light, we detected significantly more poly(A)+ RNAs cross-linked to Tm1-I/C than to the control under denaturing conditions (Fig 4B and B’), confirming the RNA binding activity of TM1-I/C. qRT-PCR of cross-linked mRNAs revealed *oskar* as a target of Tm1-I/C (Fig 4C). To identify the region of *oskar* mRNA to which Tm1-I/C binds, we incubated embryonic lysates expressing EmGFP-Tm1-I with exogenous digoxygenin-labelled *oskar* RNA fragments and subjected them to UV cross-linking. Immunoprecipitation allowed the recovery of the 3’UTR, but not of other regions of *oskar* mRNA (Fig 4D and E). Truncated (Fig S4A and B) and non-dimerizing *oskar* 3’UTR (Fig 4F) bound to EmGFP-Tm1-I with greatly reduced affinity. These findings indicate that Tm1-I/C is and RNA binding protein and that its efficient binding to *oskar* mRNPs requires an intact, dimerizing *oskar* 3’UTR.

To test whether Tm1-I/C and Khc co-exist in *oskar* mRNP complexes, we performed *oskar in situ* hybridization on EmGFP-Tm1-I-rescued *Tm1*^*gs1*^ egg-chambers carrying one copy of the *Khc*^*mKate2*^ allele (Fig 5A-C’). We found that only small portions of *oskar* mRNPs co-localized with Khc-mKate2 (~4.6%) or EmGFP-Tm1-I (~5.7%) in oocytes *in situ* (Fig 4D), similar to what we observed in our *ex vivo* co-localization analysis (Fig 2C). Interestingly, the portion of *oskar* mRNPs positive for both Khc-mKate2 and EmGFP-Tm1-I (~5.1%) was almost 40% higher than the value expected from the amount of colocalization of the mRNA with each component alone (p=10^−4^, Fig 5F, Fig S4A). This positive correlation between the presence of Tm1-I/C and Khc in *oskar* transport particles indicates that in most cases when one of the molecules is part of an *oskar* mRNPs the other molecule is present as well. In a similar analysis, we found that the EJC component Mago, although part of *oskar* mRNPs, did not exhibit such a positive correlation of colocalization with Tm1-I on the mRNA (Fig S5A and B)

**Figure 5:**
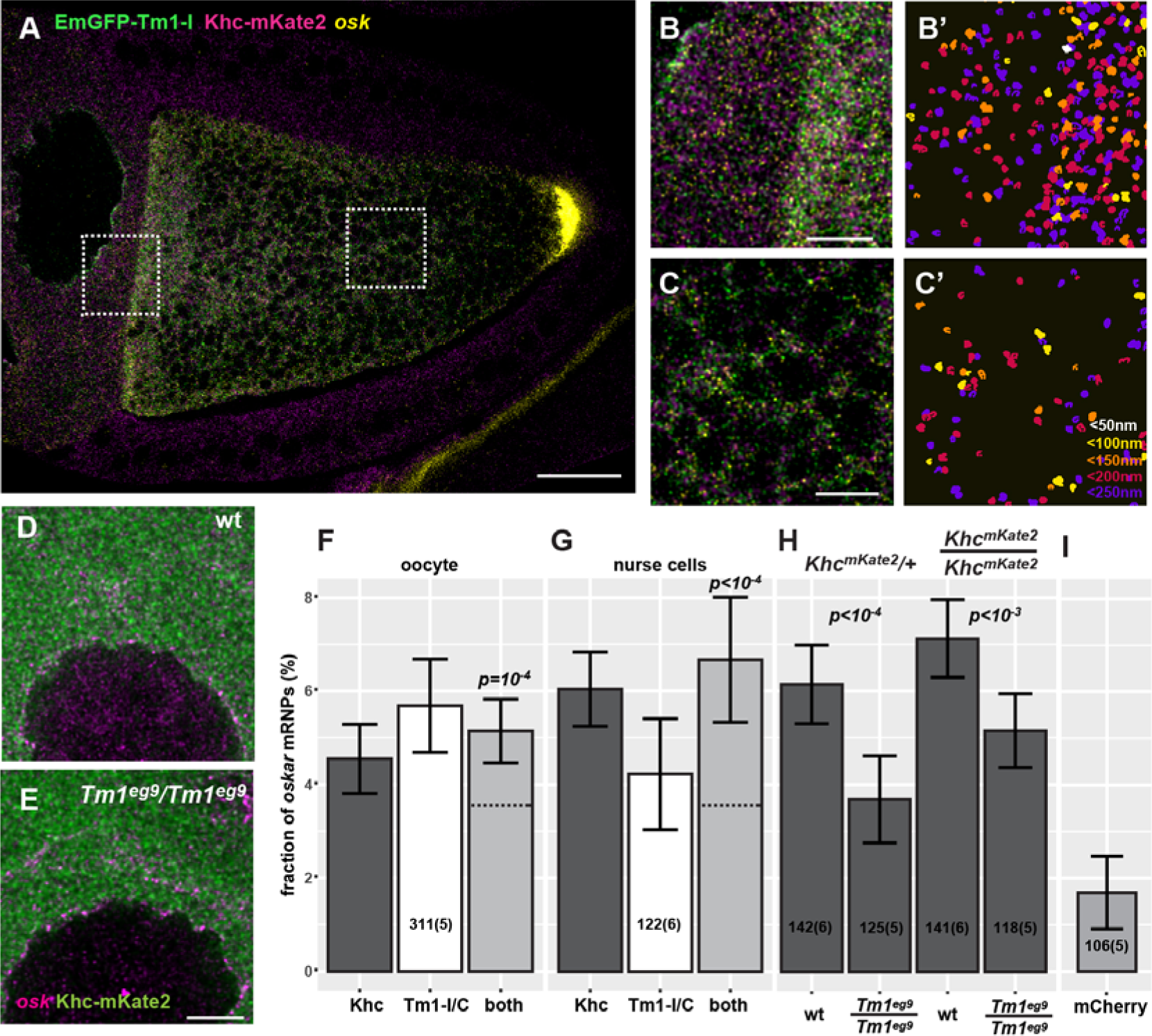
Composition of *oskar* mRNPs *in situ*. (A, B, C) Confocal image of a *Tm1*^*eg9*^ homozygous egg-chamber expressing EmGFP-Tm1-I (green) and Khc-mKate2 (magenta). *oskar* mRNA labelled with osk1-5 FIT probes(Hovelmann, Gaspar et al., 2014) is in yellow. (B’, C’) *oskar* mRNPs co-localizing with both EmGFP-Tm1-I and Khc-mKate2. Colors indicate the maximal co-localization distance (C’). Panels B-C’ represent the boxed regions in panel A. (D, E) Localization of Khc-mKate2 (green) and *oskar* mRNA (magenta) in wild-type and *Tm1*^*gs*^ mutant nurse cells. (F, G) Fraction of *oskar* mRNPs co-localizing with Khc-mKate (dark grey), EmGFP-Tm1-I (white), or both of these proteins (light grey) in the oocyte (F) or in the nurse cells (G) (max. colocalization distance is 250 nm). None of the values are significantly different from each other (one-way ANOVA, p>10^−3^). Horizontal dashed lines indicate the expected value of observing both protein in an *oskar* mRNP if the interactions are independent (see Fig S4C and D). Significance of the observed co-localization values versus the expected values are shown. (H) Fraction of *oskar* mRNPs co-localizing with Khc-mKate in wild-type and *Tm1*^*gs*^ mutant nurse cells when half or all Khc molecules are labelled (as indicated above the graph). P values of pair-wise t-tests are indicated. (I) Fraction of *oskar* mRNPs co-localizing with free mCherry in wild-type nurse cells used as negative control. The measured fraction (~1.6%) is significantly different from zero (one sample t-test). All other measured co-localization values are significantly different form this negative control (p<0.001, one-way ANOVA). Numbers indicate the number of particle clusters (100 *oskar* mRNPs in each) and the number of egg-chambers (in brackets) analysed. Error bars represent 95% confidence intervals. All values (F-I) are significantly different from zero (p<10^−3^, one sample t-test). Scale bars represent 20 µm (A) 5 µm (B, C, E). See also Figs S2 and S4.

## Kinesin-1 is recruited by Tm1-I/C to *oskar* upon nuclear export

In the course of our FISH co-localization analysis, we noted that Khc-mKate2 and EmGFP-Tm1-I colocalized with *oskar* mRNPs not only in the oocyte, but also in the nurse cell cytoplasm (Fig 5A-B’ and G). To test whether this colocalization of Khc with *oskar* mRNA also requires Tm1-I/C, we analysed Khc-mKate2 association with endogenous *oskar* mRNA in *Tm1*^*gs*^ mutant nurse cells (Fig 5D and E). We found an almost two fold reduction in Khc positive *oskar* mRNPs in absence of Tm1-I/C when a single *Khc* allele was labelled (Fig 5H). When all Khc molecules were fluorescently tagged, we also detected a significant difference in Khc association in *Tm1*^*gs*^ mutant and wild-type nurse cells, although we did not observe the almost two-fold increase in Khc positive *oskar* mRNPs in the wild-type controls that we observed in case of the *Tm1*^*gs*^ mutant (Fig 5H). This observation highlights the possible limitation of our co-localization analysis when crowding of at least one of the objects occurs (Fig 5D and E). Although STED superresolution microscopy further increased crowding by resolving the confocal objects (Fig S5C, C’ and G), it confirmed that both Khc-EGFP or EmGFP-Tm1-I are recruited to *oskar* mRNPs (Fig S5D). Moreover, it reinforced our finding that Khc-EGFP association to *oskar* mRNPs in the nurse cells is greatly reduced in the absence of Tm1-I/C (Fig S5E).

Fluorescently tagged Tm1-I, similarly to Khc-mKate, localized diffusely in the cytoplasm and, unlike other tropomyosins, did not accumulate on actin structures in the egg-chamber (Fig 6C-D’). In contrast, Tm1-I accumulated at the posterior pole of the oocyte (Cho, Kato et al., 2016) (Fig S6A, C’, E-G). Interestingly, we also detected the fluorescent Tm1-I signal in the nurse cell nuclei (Fig 6A, Figs S5F and 6B), however, in contrast to GFP-Mago, we did not find evidence that nuclear Tm1-I/C associates with *oskar* transcripts (Fig S5H).

**Figure 6:**
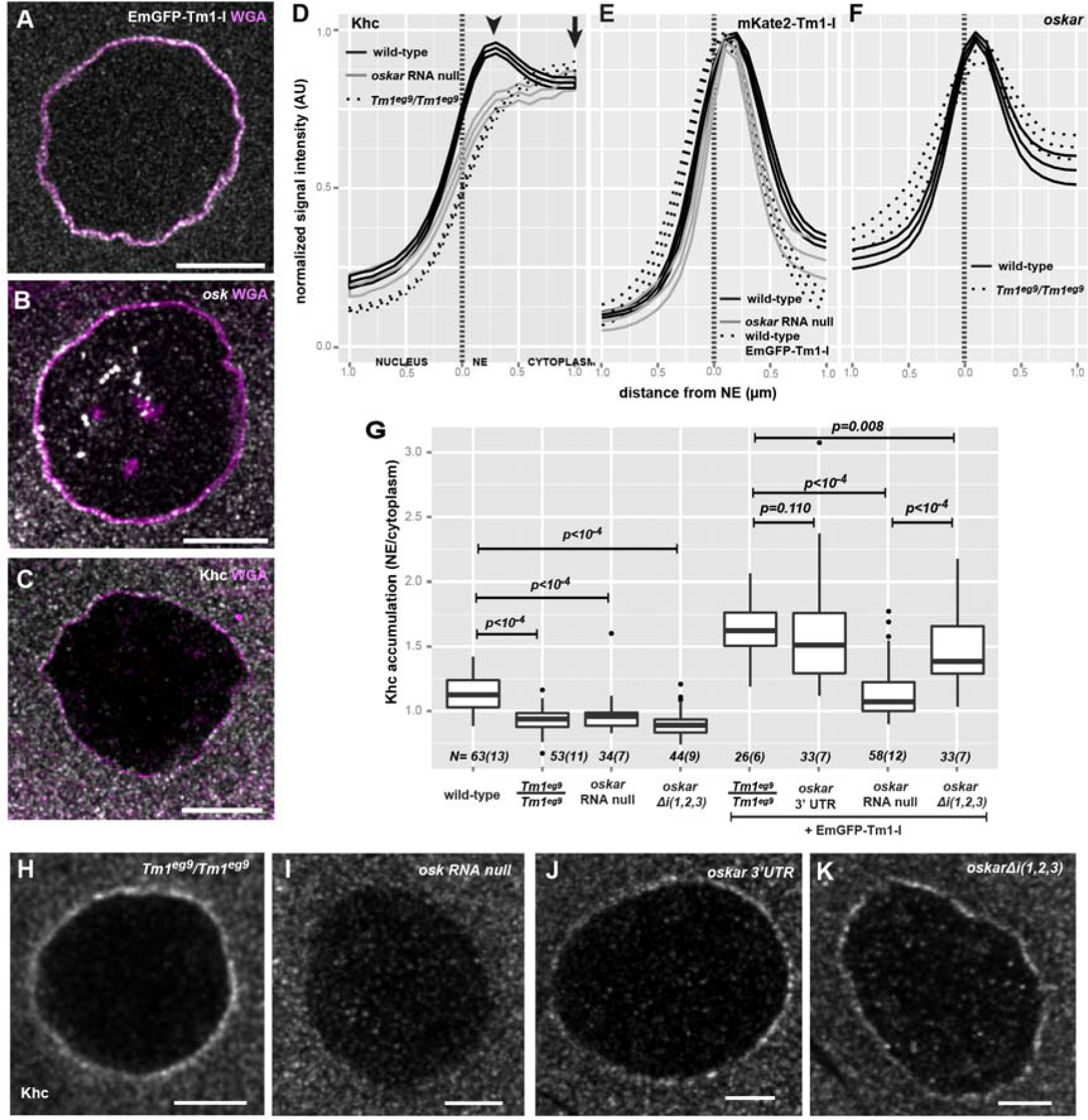
Transient accumulation of *oskar* mRNPs around the NE. (A-C) Localization of EmGFP-Tm1-I (A) *oskar* mRNA (B) and Khc (C) around the nurse cell nuclear envelope (magenta, WGA staining). (D-F) Mean distribution profile of Khc (D) mKate2-Tm1-I, EmGFP-Tm1-I (E) and *oskar* mRNA (F) around the NE of nurse cells (marked with a dotted vertical line). Genotypes are indicated as follows: wild-type control with solid black line (D-F), *oskar* RNA null with solid grey line (D, E), *Tm1*^*eg9*^/*Tm1*^*eg9*^ with dotted black line (D, F) Lines indicate mean±95% conf. int. We analyzed the accumulation of mKate2-Tm1-I instead of EmGFP-Tm1-I due to unexpected GFP expression from the *oskar*^*0*^ allele. (G) Khc accumulation around the NE. To calculate accumulation the signal intensity measured at the position of the peak observed in the wild-type control (D, arrowhead, 356±17.6 nm away from NE) was divided by the signal intensity 2 *s*.d. away (d, arrow, at 356+2*410 nm). P values of pairwise Mann-Whitney U tests against wild-type control or *Tm1*^*gs*^ rescued with EmGFP-Tm1-I are indicated above the boxplots. Numbers indicate the number of nuclei and the number of egg-chambers (in brackets) analysed. (H-K) Khc accumulation around the NE of nurse cells over-expressing EmGFP-Tm1-I within *Tm1*^*eg9*^/*Tm1*^*eg9*^ (H), and *oskar* RNA null egg-chambers (I) expressing either the *oskar* 3′ UTR (J) or *oskar Δ*_*i*_*(1,2,3)* (K). Scale bars are 10 µm (A-C), 5 µm (H-K)

The most prominent localization pattern of the fluorescently tagged Tm1-I was its enrichment around the nuclear envelope (NE) in the nurse cells (Fig 6A, Fig S6) (Cho et al., 2016, Veeranan-Karmegam et al., 2016), similar to that of *oskar* mRNA (Little, Sinsimer et al., 2015) (Fig 6B). We also detected perinuclear enrichment of endogenous Tm1-I/C on immunolabeled wild-type specimens (Fig S6G and I), although it was less pronounced due to the high aspecific signal in the nurse cell cytoplasm (Fig S6H). We observed that Khc also enriches around the nurse cell NE (Fig 6C), although to a lesser extent than the fluorescently tagged Tm1-I/C or *oskar* mRNA detected by FISH. Our analysis of radial profiles of NEs counterstained by fluorescent lectins, confirmed a slight but significant perinuclear accumulation of Khc in wild-type nurse cells (Fig 6D and G), independent of the developmental age of the egg-chamber (Fig S7A). This observed Khc accumulation around the NE required the presence of both *oskar* mRNA and of Tm1-I/C (Fig 6D and G). In contrast, *oskar* mRNA or mKate2-Tm1-I accumulation around the NE was not affected in *Tm1*^*gs1*^ or *oskar*-null mutant egg-chambers, respectively (Fig 6E and F).

We noted that transgenic over-expression of EmGFP-Tm1-I increased Khc recruitment to the NE substantially in the rescued *Tm1*^*gs*^ egg-chambers (Fig 6G and H). Although there was a slight elevation in Khc accumulation in the absence of *oskar* RNA (Fig 6G and I), possibly reflecting the presence of other, even non-specific mRNA targets of over-expressed EmGFP-Tm1-I, the substantial increase was only observed when an intact *oskar* 3’UTR, whether in endogenous *oskar* mRNA, transgenic *oskar Δ*_*i*_*(1,2,3)* or the *oskar* 3’UTR alone, was present (Fig 6G, J and K). Given that even in the absence of *oskar* mKate2-Tm1-I was enriched around the NE, these results not only confirm the instrumental role of Tm1-I/C in the Khc recruitment process, but also indicate that kinesin-1 loading on *oskar* mRNPs takes place if and only if mRNAs containing the *oskar* 3’UTR are available. Consistent with this, in the nurse cell cytoplasm of *oskar* RNA null egg-chambers expressing the *oskar* 3’UTR and Khc-mKate2, we detected an identical degree of Khc association with *oskar* mRNPs as observed in wild-type control egg-chambers (Fig S5I, J).

### Kinesin recruited by Tm1-I/C is inactive

To address the functional consequences of the ‘super-loading’ of Khc at the NE upon EmGFP-Tm1-I over-expression, we quantified the mean distribution of the non-localizing *oskar Δi(1,2,3)* RNA throughout stage 9 oocytes (Gaspar et al., 2014). This analysis showed that over-expression of EmGFP-Tm1-I causes a substantial posterior-ward shift of the non-spliced *oskar* mRNA (Fig 7A and B). However, the rescue of localization was not complete as it still deviated substantially from the wild-type control (Fig 7C). Furthermore, EmGFP-Tm1-I over-expression did not promote posterior localization of an RNA consisting solely of the *oskar* 3’UTR (Fig 7D), although the posterior enrichment of Khc was not affected (Fig S7F and G). EmGFP-Tm1-I over-expression also did not promote *oskar* mRNA localization in oocytes with reduced Khc levels (Fig 7E, Fig S7H), confirming the essential role of kinesin-1 motor is this process. These observations indicate that a properly assembled EJC/SOLE is required to activate the *oskar* 3’ UTR bound, Tm1-I/C recruited kinesin-1 within the oocyte during mid-oogenesis.

**Figure 7:**
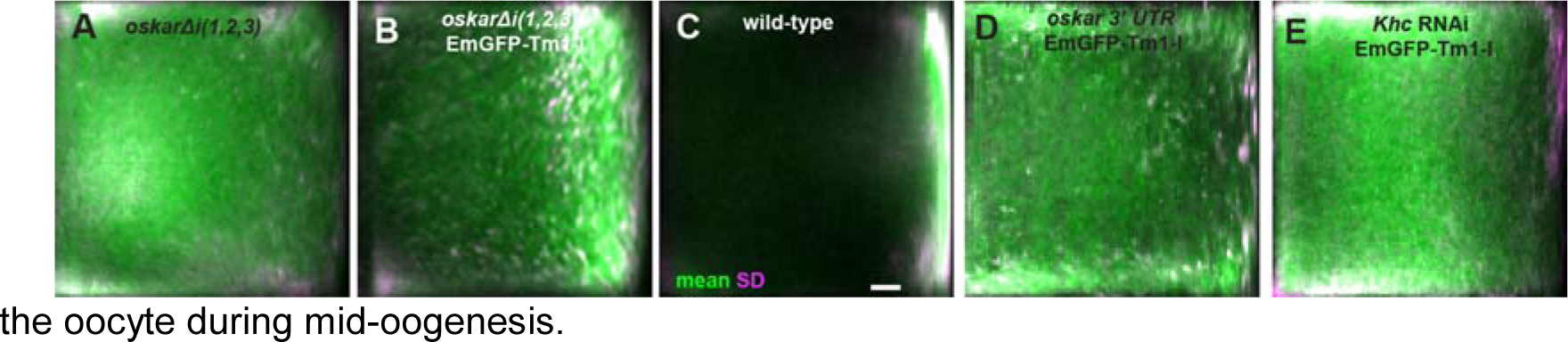
Effects of EmGFP-Tm1-I overexpression on *oskar* mRNA localization. (A-E) Mean *oskar* mRNA distribution (green, detected by conventional FISH) within oocytes in which *oskar* mRNA is substituted by *oskar Δi(1,2,3)* mRNA (A, B), *oskar* 3’UTR mRNA (D) and that (in addition) over-express EmGFP-Tm1-I (B, D and E). Wild-type control (C) and oocytes expressing *Khc* RNAi and EmGFP-Tm1-I (E). Magenta indicates standard deviation of the distribution of *oskar* mRNA. Scale bar is 10% of total oocyte length (C). See also Fig S7.

## Discussion

Loading of the appropriate transport machinery on mRNPs is critical to achieve correct localization and, consequently, localized translation of the transcript. Although mRNA transport has been extensively studied over the last two decades, the recruitment of plus-end directed kinesin motors to mRNPs and their regulation remains poorly understood (Medioni et al., 2012).

Here, we have shown that the majority of kinesin-1 motor associated with *oskar* mRNA is recruited by Tropomyosin1-I/C, a non-canonical RNA binding protein. We found that this recruitment occurs early in the cytoplasmic life of the mRNA, upon its nucleo-cytoplasmic export, in the perinuclear cytoplasm of the nurse cells, where dimerization of *oskar* mRNA molecules via their 3’UTRs commences (Little et al., 2015). Our observations are consistent with an “ergonomic” kinesin-loading machinery that becomes functional when and only when Tm1-I/C binds the *oskar* 3’ UTR in the nurse cell cytoplasm. The kinesin-recruitment machinery is inefficient, as only a small portion of *oskar* mRNPs are kinesin-bound even during the most active phase of *oskar* mRNA posterior-ward transport in the oocyte. Such inefficiency may serve to prevent the sequestration of kinesin-1 molecules by *oskar* mRNPs away from other cargoes requiring the motor for transport. On the other hand, the transient and dynamic binding and unbinding of the motor may enable transport of virtually all *oskar* mRNPs in a temporally coordinated fashion, thereby promoting localization of more than 50% of *oskar* mRNA to the posterior pole of the oocyte by the end of stage 9 (Gaspar et al., 2014). Additionally, as *oskar* mRNA is continuously transported from the nurse cells to the oocyte from the beginning of oogenesis (Ephrussi & Lehmann, 1992, Jambor et al., 2014, Kim-Ha et al., 1991), the age of *oskar* mRNPs at the onset of posterior-ward transport may vary between a few minutes to one-one and a half days. The dynamic recruitment of Khc may also guarantee that the *oskar* particles are equipped by transport competent motor molecules at any moment.

The first step of *oskar* transport is mediated by cytoplasmic dynein (Clark et al., 2007, Jambor et al., 2014), which is presumably also recruited at the nurse cell NE, where we detected the accumulation of the RNA cargo adapter Egalitarian (Dienstbier et al., 2009, Navarro, Puthalakath et al., 2004) and the dynactin component Dynamitin (McGrail, Gepner et al., 1995). Interestingly, the dynein apoenzyme did not enrich around the NE (Fig S7B-E), possibly because its association with *oskar* mRNPs instantly initiates their transport away from the NE into the oocyte. Within the oocyte, efficient posterior-ward transport of *oskar* mRNA only takes place upon repolarization of the MT cytoskeleton during mid-oogenesis (Parton, Hamilton et al., 2011, Theurkauf, 1994). Therefore the activity of the kinesin-1 recruited to *oskar* mRNPs must be regulated and co-ordinated with that of the dynein in response either to environmental changes (Burn, Shimada et al., 2015, Gaspar et al., 2014) or to the developmental program. Our data showing the failed or incomplete posterior localization of *oskar* 3’UTR and *oskar Δi(1,2,3)*, respectively, indicates that the kinesin-1 recruited to the *oskar* 3’UTR by Tm1-I/C is inactive. Since the *oskar* coding sequence – with the exception of the SOLE – is dispensable for the localization process (Ghosh et al., 2012), we propose that, during mid-oogenesis, the spliced, EJC associated SOLE complex activates the *oskar* RNA-bound kinesin-1. Therefore, although the EJC/SOLE is necessary for proper *oskar* mRNA localization, it not sufficient, as recruitment of the kinesin-1 motor to *oskar* mRNPs is mediated Tm1-I/C, whose RNA binding scaffold is provided by the *oskar* 3’ UTR. To our knowledge, *oskar* is the first mRNA and the first cargo of kinesin-1 described where loading and activation of the motor are decoupled and an unproductive tug-of-war between two opposing motors is avoided by keeping one of the molecules in a long-term stasis.

Although the underlying mechanisms of the Khc loading and activation processes remain cryptic, a recent study demonstrated that the deletion of the ATP-independent MT binding site and the auto-inhibitory IAK motif of Khc results in phenotypes consistent with a failure in motor activation and/or recruitment to *oskar* mRNPs (Williams, Ganguly et al., 2014).

Moreover, it was shown parallel with our study that Tm1-I/C is able to directly bind Khc *in vitro*, and that this interaction depends on the ATP-independent MT-binding site (Veeranan-Karmegam et al., 2016), indicating that Tm1-I/C directly links kinesin-1 to *oskar* mRNA. This unconventional mode of cargo binding via the unconventional cargo adapter Tm1-I/C may result in incomplete release of kinesin from auto-inhibition and thus explain the persistent inactivity of the kinesin-1 upon its recruitment to the mRNA.

Tm1-I/C is an atypical tropomyosin: it does not enrich around microfilaments in the female germ-line (Cho et al., 2016, Veeranan-Karmegam et al., 2016, and this study), forms intermediate filament like structures *in vitro* (Cho et al., 2016), and directly binds to Khc (Veeranan-Karmegam et al., 2016) and to RNA, recruiting kinesin-1 dynamically to *oskar* mRNPs. Although Tm1-I/C has a short tropomyosin superfamily domain in its C-terminal moiety, most of the protein is composed of low complexity sequences (Cho et al., 2016). Such intrinsically disordered proteins (IDPs) are often constituents of RNA containing membraneless organelles, such as RNA granules (Kato, Han et al., 2012), stress granules (Molliex, Temirov et al., 2015), P granules (Elbaum-Garfinkle, Kim et al., 2015), nuage and germ granules(Nott, Petsalaki et al., 2015). Furthermore, it has been shown that IDPs fold upon binding to their partners (Wright & Dyson, 2009). This may explain why Tm1-I/C only recruits Khc to the NE in the presence of the *oskar* 3’UTR. Further dissection of the structure, the precise molecular functions of Tm1-I/C, its binding partners will be crucial to determining the nature and the minimal number of features necessary for such unconventional yet vital kinesin-1 mediated localization of mRNA.

## Methods and Materials

### Fly stocks

*Tm1*^*eg1*^ and *Tm1*^*eg9*^ (FBal0049223) mutations originating from the imprecise excision of the P-element in *Tm1*^*gs1*^ (FBal0049228) were used to study the role of Tm1 in *oskar* mRNA localization (Erdelyi et al., 1995). To remove *oskar* mRNA (*oskar* RNA null), a newly created *osk*^*attP,3P3GFP*^ (*oskar*^*0*^ in this manuscript, see below) allele was used in homozygous form or in combination with another RNA null allele, *osk*^*A87*^ (FBal0141009). We observed no difference between the phenotype of *osk*^*attP,3P3GFP*^ homo-and the *osk*^*attP,3P3GFP*^/*osk*^*A87*^ heterozygotes in our assays. The following *oskar* mRNA mutations and truncations were expressed in an *oskar* RNA null background: *UASp-osk.3’UTR* (Filardo & Ephrussi, 2003) (FBal0143291) *UASp-oskar Δi(1,2,3) 5x BoxB* (Ghosh et al., 2012) (no protein coding function, FBal0291667). *Tub67C::GFP*^*m6*^-*Staufen* (Schuldt, Adams et al., 1998) (FBal0091177), *UASp-dmn-GFP* (Januschke, Gervais et al., 2002) (FBal0145074) α*Tub::Khc-EGFP* (Sung et al., 2008) (FBal0230204) *UASp-GFP-Mago (Newmark, Mohr et al., 1997)* (FBal0063884) and Ketel-GFP (Villanyi, Debec et al., 2008) (FBal0244142) were used to visualise Staufen, Dynamitin, Khc, Mago and Importin-β molecules. To express mCherry FP in the nurse cells, we used the TM3, P{sChFP} balancer chromosome (FBti0141181). The *Khc*^*27*^ protein null allele (FBal0101625) was used to halve Khc levels (heterozygous) or completely remove Khc from egg-chambers developing from *Khc*^*27*^ homozygous germ-line clones, *UASp-Khc RNAi* Trip Line GL00330 (Staller, Yan et al., 2013) was used to knock down Khc levels. To label *oskar* mRNPs we used the *osk::oskMS2(10x)* system together with *hsp83::MCP-EGFP* (Zimyanin et al., 2008) or *hsp83::MCP-mCherry* (gift of L. Gavis) and *UASp-EB1-mCherry* (gift of D. Brunner) to label the growing plus ends of MTs. For the Khc tethering experiment, we expressed *osk::oskMS2(6x)* (Lin, Jiao et al., 2008) (FBal0263509), as *oskMS2(10x)* is not translated at the posterior pole (Zimyanin et al., 2008) Expression of all UASp transgenic constructs were driven with one copy of *oskar-Gal4* (Telley, Gaspar et al., 2012) (FBtp0083699) with the exception of the *Khc* RNAi line, where a second *oskar-Gal4* allele was introduced to boost the expression level of co-expressed *UASp-EmGFP-Tm1-I*. *w*^*1118*^ (FBal0018186) was used as wild-type control. All stocks were raised on normal cornmeal agar at 25 °C.

### Transgenic constructs and new alleles

*UASp-EmGFP-Tm1-I* and *UASp-mKate2-Tm1-I* were created by fusing the EmeraldGFP and the mKate2coding sequences in frame with the 5’ of the Tm1-I coding sequence (cds), respectively, followed by the 3′ UTR amplified from an ovarian cDNA. These construct were completed with the 5’ UTR of Tm1-I and inserted into pUASp2 transgenesis vectors (Rorth, 1998).

*UASp-Khc*_*401*_*-MCP* was cloned by ligating the sequence encoding the N terminal-most 401 aminoacids of Khc -sufficient for processive movement (Telley et al., 2009) – with the cds of MS2-coat protein (MCP) in frame in a pUASp2 transgenesis vector.

The *oskar* RNA null allele, *oskar*^*attP,3P3GFP*^ (*oskar*^*0*^) was created by homologous recombination mediated knock-in. A double-stranded break in the *oskar* promoter 26 nucleotides upstream of the transcriptional start site was induced by CRISPR/Cas9, by co-injecting a Cas9 expressing vector, a U6 promoter driven guide RNA (5’GAATGGAGAAGTGGACCGCGT**NGG**) and a template plasmid carrying a minimal attB-loxP-GMR-3P3-EGFP-tubulin3’UTR-loxP cassette flanked by two ~1000 bp long homology arms (Sebo, Lee et al., 2014). Female flies homozygous for *oskar*^*attP,3P3GFP*^ are sterile, oogenesis is terminated prematurely at around stage 6, and oocytes express the same karyosome defects as previously reported by Jenny et al. (Jenny, Hachet et al., 2006). The *oskar* promoter drives EGFP expression from the *3P3-EGFP* marker gene in the female germline which may interfere with the read-out of another GFP tagged transgene. Cre mediated removal of the *3P3-EGFP* marker gene restores female fertility.

The genomic Tm1 locus was tagged with the *mCherry* cds (*Tm1*^*mCherry*^) using a similar strategy as described for *oskar*^*attP,3P3GFP*^, using the guide RNA cds (5′GGAGTCATCGTGCTCCATGT**NGG**) and a loxP-GMR-3P3-EGFP-tubulin3’UTR-loxP-mCherry cassette flanked by two ~ 600 bp long homology arms targeting the 5’UTR/ATG boundary to insert the *mCherry* sequence in frame with Tm1-I.

Expression of mCherry-Tm1 was achieved by Cre mediated stable removal of the *3P3-GFP* marker gene from the locus. Flies homozygous for *Tm1*^*mCherry*^ are viable, however have reduced life span and are flightless, probably due to internally tagged and thus compromised Tm1-E and Tm1-H isoforms. Both homozygous males and females are fertile.

The genomic Khc locus was tagged with the *mKate2* cds (*Khc*^*mKate2*^) using a similar strategy as described above. The genome was targeted using a guide RNA cds (5 GATTGGATCTACGAGTTGAC**NGG**) and an mKate2 cassette flanked by two ~ 700 bp long homology arms targeting the STOP/3’UTR boundary to insert the *mKate2* sequence in frame with Khc. F1 generation embryos were screened for the mKate2 fluorescence to identify individuals with modified genome. Flies homozygous for *Khc*^*mKate2*^ are viable and fertile.

### Immunological techniques

Immunoprecipitation of EmGFP-Tm1-I and Flag-Myc-GFP (as control) from ovarian lysates was carried out as described in Ghosh et al(Ghosh, Obrdlik et al., 2014). Eluates were tested in western blot analysis probing with anti-Khc (1:5000, Cytoskeleton), anti-Staufen (1:1000)(Krauss, Lopez de

Quinto et al., 2009), anti-Bruno (1:5000)(Chekulaeva, Hentze et al., 2006), anti-Y14 (1:2500)(Hachet & Ephrussi, 2001), anti-BicD (2:100, DHSB clone #1B11 and 4C2), anti-Dic (1:2000, Millipore) and anti-GFP (1:2000, Millipore). The same antibodies and dilutions were used to detect RNAi knock-down efficiency and relative levels of Khc-EGFP. To test Tm1-I/C expression in ovarian lysates a pan-Tm1 antibody (1:1000)(Cho et al., 2016) recognizing the Tropomyosin domain shared by the Tm1 isofoms was used (kind gift of D. Montell).

Khc, Tm1, Dic and Egl were visualized in heat fixed egg-chambers 23 incubated overnight at 4 °C in anti-Khc (1:250), pan-Tm1 (1:500), anti-Dic (1:1000) or anti-Egl (1:1000) primary antibodies diluted in PBST (PBS+0.1%Triton-X-100)/10% normal goat serum. Signal was developed by applying AlexaFluor488 or Cy5 conjugated anti-rabbit or anti-mouse secondary antibodies for 60 minutes at room temperature (RT) (1:1000, Jackson ImmunoResearch). GFP-autofluorescence was preserved by fixation for 20 minutes in 2%PFA/0.05%Triton-X 100 in PBS. To visualize F-actin, samples were fixed in 2% PFA/0.05% Triton-X 100 in BRB80 (80mM PIPES, 2 mM MgCl_2_, 1mM EGTA) and stained with phalloidin-FITC (1:100, Molecular Probes) and all washes were carried out in BRB80+0.1%

Triton-X 100.

NE was counterstained by WGA-TRITC (1:300, Life Sciences) for 60 minutes at RT. Samples were embedded in 80%glycerol+2% N-propyl-gallate mounting medium.

### RNA recovery after cross-linking

To test if Tm1-I/C binds RNA directly, 0-2h old embryos expressing EmGFP-Tm1-I, EGFP or FlagMycGFP were collected and 1 J/cm2 UV (*λ* = 254 nm) was applied to cross-link nucleic acids with protein molecules at zero distance. Embryos were lysed in 10x RIPA buffer composed of low salt buffer (20 mM Tris-CL, 150 mM KCL, 0.5 mM EDTA, pH=7.5) containing strong detergents (1% Triton-X100, 1% deoxycholate and 0.1% SDS), reducing agent (5mM DTT) and 5x protease inhibitor cocktail (PIC, Roche), crude debris was removed by 10 minute long centrifugation with 16000 g at 4 °C. Clarified lysates were diluted to 10mg/ml with 10x RIPA buffer, then further diluted ten-fold with low salt buffer. 1 mg protein of lysate (1ml) was supplemented with 10 µl of GFP-Trap_M beads (Chromotek) and immunoprecipitation was carried out at 4 °C for 90 minutes. Beads were washed six times with high salt buffer (20 mM Tris-CL, 500 mM KCL, 1 mM EDTA, 0.1% SDS, 0.5 mM DTT, 1x PIC pH=7.5) at room temperature for a total duration of 60 minutes (stringent washes). Alternatively, beads were washed twice with high salt buffer then three times under denaturing conditions with 7.6 M urea and 1%SDS in PBS (Chromotek Application Note). RNA was then extracted from the washed beads for qRT-PCR or FISH was performed on the beads with oligo(dT)25-TexasRed similarly as described in(Castello, Fischer et al., 2012). FISH-on-beads experiments were imaged on a Leica SP8 with a 63x 1.4 NA oil immersion objective. Bead contours were captured by imaging the reflection of 594 nm light, oligo(dT)25 probes were excited with 594 nm. Primer sequences used during qRT-PCR are available upon request.

To analyse *oskar* mRNA fragments binding to EmGFP-Tm1-I, embryos were lysed in pre-XL buffer (20 mM Tris-CL, 150 mM KCL, 4mM MgCl2, 2x PIC, pH=7.5), cleared from debris and lysate concentration was adjusted to 10 mg/ml. 0.2-0.8 ng/nucleotide DIG labelled RNA (1:10 ratio of labelled to unlabelled uracil) and 1µl RiboLock (ThermoFisher Scientific) were added to 100 µl lysate. Mixtures were incubated for 20 minutes at RT prior to cross-linking with 254 nm UV light 0.15 J/cm2 energy. Subsequently, the lysate was diluted with an equal volume of low salt buffer supplemented with detergents, reducing agent and PIC (see above) and further diluted 1:5 with low salt buffer. 2 µl RiboLock and 5 µl GFP-Trap_M beads were added per 500 µl volume, and immunoprecipitation and washes were carried out as described above. Beads were then labelled with anti-GFP-CF488A (1:1000, Sigma Aldrich) and anti-DIG-Hrp (1:500, Roche) PBST for 30 minutes. Anti-DIG signal was developed using Cy5-TSA amplification kit (Perkin Elmer). Fluorescently labelled beads were mounted on slides in glycerol based mounting medium and were imaged with a Leica 7000 TIRF microscope using a 100x 1.46 NA oil immersion objective, 1.6x Optovar and epifluorescent illumination.

### Fluorescent *in situ* hybridization

For conventional FISH, specimens were fixed in 2%PFA/0.05%Triton-X 100 in PBS for two hours at RT. After fixation and washes in PBST, 2 µg/ml Proteinase-K was applied for 5 minutes at RT, followed by 5 minutes boiling at 92 °C in PBS/0.05% SDS. The samples were then pre-hybridized for 60 minutes at 65 °C in hybridization buffer (HYBEC: 5x SSC, 15% ethylene carbonate, 50 µg/ml heparin, 0.1mg/ml salmon sperm DNA, 0.05% SDS, pH=7.0). The *oskar* cds (targeting nucleotides 1442-1841) and *oskar* 3′ UTR (targeting 2342-2730) probes were diluted in HYBEC in 0.5 pg/nucleotide/ml concentration each. The *oskar* cds probes were directly labelled with either Atto-488 (Atto-tec) or AlexaFluor 555 (ThermoFisher Scientific), while the *oskar* 3′ UTR was directly labelled with Atto-633 (Atto-tec). Hybridization was carried out at 65 °C overnight and excess probe was removed by four washes at 65 °C (HYBEC, HYBEC/PBST 1:1, 2x PBST, each 20 minutes) and one 20 minute long wash in PBST at RT. During conventional FISH, WGA-FITC was applied to counterstain NEs. Samples were embedded in 80%glycerol+2% N-propyl-gallate mounting medium.

To preserve GFP and mKate2 auto-fluorescence, forced intercalation (FIT) probe based RNA detection was performed (Hovelmann et al., 2014). Ovaries were briefly fixed as described above, and the incubation buffer was changed to IBEX (10 mM HEPES, 125mM KCl, 1mM EDTA, 0.3% Triton-X 100, pH=7.7) after fixation. *oskar* mRNA was labelled with osk1-5 LNA modified FIT probes 45 in IBEX+ (IBEX supplemented with 15% ethylene carbonate, 50 µg/ml heparin and 10 % 10 kDa dextran sulphate) to a final concentration of 0.05 µM each. Samples were incubated at 42 °C for 30 minutes, then briefly washed in IBEX and IBEX/BRB80 (1:1 mixture) at 42 °C.

To detect *oskar* 3’ UTR with single molecule FISH (smFISH), we used 15 different 3’UTR targeting probes labelled with a single Atto-565-ddUTP nucleotide using TdT (Table S2). smFISH was carried out similarly to conventional FISH. Protease-K digestion and heat denaturation of RNA secondary structures were omitted to preserve mKate2 fluorescence. The hybridization was performed at 37 °C for 2 hours using 5nM/probe concentration.

Specimens were embed in 79% TDE (η =1.475)(Staudt, Lang et al., 2007), which boosted the brightness of GFP and mKate2 by about 2-fold and of FIT probes 4-6 fold compared to conventional glycerol based mounting media (data not shown).

### Microscopy

Conventional laser scanning confocal microscopy was carried out using a Leica TCS SP8 microscope with a 63x 1.4 NA oil immersion objective. STED microscopy was performed on a Leica STED 3x microscope with a 100x 1.4 NA oil immersion objective and HyD time-gated photodetectors. Acquired images were deconvolved with Huygens Professional (SVI) prior to analysis.

To minimize cross-talk of the two labels, GFP and TO labelled FIT probes were excited with 470 nm and 525 nm lines of a white light laser source, respectively. Emission was recorded between 480-520 nm (GFP) and 525-575 nm (TO). For similar considerations, Atto-565 and mKate2 were excited by 561nm and 610 nm light and emitted fluorescence was detected between 565-585nm and 620-720 nm, respectively. Under these conditions less than 1% of recorded signal originated from cross-talk.

To stimulate emission of the GFP and TO dyes, a 592 nm depletion donut shaped laser beam was used, with all power assigned to improve lateral resolution by about 2.5-3 fold. Typically a stack of seven slices was recorded (voxel size: 22x22x180 nm) and subsequently deconvolved with Huygens Professional. The middle slice was then subjected to object-based colocalization analysis.

### *Ex vivo* ooplasmic preparation

Crude ooplasm was obtained from living stage 9 oocytes expressing *oskMS2(10x)*,MCP-EGFP, oskGal4,EB1-mCherry for mRNP tracking or oskMS2(10x), MCP-mCherry and a GFP tagged protein of interest for *ex vivo* colocalization analysis. Ovaries were dissected in BRB80 (80mM PIPES, pH=6.9, 2mM MgCl_2_, 1mM EGTA). BRB80 was replaced with 1% IB (10 mM HEPES, pH=7.7, 100 mM KCl, 1 mM MgCl_2_, 1% 10kDa dextran) and ovaries were transferred onto silanized coverslips. Silanization was carried out with dichlorodimethylsilane (Sigma-Aldrich) under vacuum for 1.5-2h to obtain a slightly hydrophobic surface. A drop of Voltalef 10S oil (VWR) was placed next to the dissected ovaries and individual ovarioles containing stage 9 egg-chambers were pulled under oil with fine tungsten needles. There, the stage 9 egg-chambers were isolated and the nurse cell compartment was carefully removed with needles. Using a gentle pulling force at the posterior pole of the created “oocyte sack” (the oocyte and surrounding follicle cells), the ooplasm was slowly released from anterior to posterior onto the coverslip surface. The level of surface hydrophobicity was critical: a hydrophilic surface bound *oskar* mRNPs aspecifically, blocking their motility, whereas it was impossible to create an ooplasmic streak on coverslips that were too hydrophobic. Such preparations were imaged on a Leica 7000 TIRF microscope with a 100x 1.46 NA oil objective and 1.6x Optovar. Images were collected simultaneously for 32 s with a Photometrics Evolve 512 EM CCD camera with 140 nm lateral resolution. We observed no decline in RNP motility and MT dynamics within the first 60 minutes (data not shown).

### Image analysis

*In vivo* tracking of *oskMS2*-GFP particles and *oskar* mRNA distribution within oocytes was performed as previously described (Gaspar et al., 2014). Image segmentation for tracking, colocalization analysis of *oskar* mRNPs, and measurement of bead fluorescence was carried out using a custom particle detector library in ImageJ.

#### Extraction of NE radial profiles

A section containing a close to maximal cross-section of the nucleus was selected, the outline of the NE counterstained with WGA was coarsely traced manually. At each point along the outline, the signal under a 5 µm long segment perpendicular to the outline was extracted, resulting in a few hundred to thousand, roughly 2.5 µm long reads on both the cytoplasmic and the nuclear side of the outline. At each point, the position of the NE was determined with sub-pixel precision though fitting a Gaussian function to the WGA signal. All other recorded signal was positioned relative to the location of the NE and was averaged for a given nucleus. These mean signal intensities were then normalized to the maximum value of a radial profile.

#### Ex vivo

tracking was done automatically, tracks displaying linear displacements were manually selected and their directionally manually assigned by overlaying them with the EB1 channel. 8-20% of detected tracks could not be assigned a polarity due to the absence of nearby co-axial EB1 comets (Fig S1B). Linear runs were extracted from the detected tracks using a series of custom Excel macros (Gaspar et al., 2014). Runs of unknown polarity were on average shorter than classified minus and plus end directed runs (Fig S1D).

#### Object-based colocalization

was assayed by measuring the distance between closest-neighbour objects from the *oskar* mRNP (reference) and GFP/mKate2 (target) channels within a confined area representing exclusively the nurse cell perinuclear region and cytoplasm. Random colocalization was addressed by seeding the objects of the target channel randomly into the confined area. This simulation was repeated one hundred times to obtain a distribution of expected (random) colocalization. To calculate fraction of colocalization and to compensate for the huge variability of observed particles per image, reference channel objects were randomly assigned into particle clusters representing 160 particles for *ex vivo* and *in situ* colocalization analysis, and 100 particles for competitive FISH. With these values, only ~5% of observed objects of the reference channel were excluded from the analysis. Observed colocalization within each cluster was compared to the distribution of simulated random values using one sample Student’s t-test (α = 0.01). Significant values were used to calculate the difference between observed and random colocalization to assess true, biological colocalization. These differences were found to be significantly different from zero for all analysed protein molecules in wild-type samples, except for the negative control Ketel-GFP (Fig. S5D). By opening the colocalization window (the maximal inter-neighbour distance), random colocalization rapidly overcomes the observed values (e.g. Fig S1E-H) due to particle crowdedness both *ex vivo* and *in situ* resulting in a probable underestimation of true colocalization. To minimize this effect, the clipping point where the difference was maximal was determined, and the halfway distance between zero and the clipping point was used to compare the effects of different conditions on *oskar* mRNP composition. This colocalization window was 200 nm *ex vivo* (non-fixed specimen), 250 nm *in situ* and 100 nm STED *in situ* (fixed).

#### Temporal co-localization

was assayed as described in the legend of Fig S3C. Importantly, the same microscope settings were used to acquire image sequences of a given protein molecule (e.g. Khc-mKate2) that allows direct comparison of signal intensities between hetero-and homozygous extracts (see thresholding in the legend of Fig S3C).

Transformations and statistical analysis of all the obtained numerical data were carried out in R (Team, 2012) using the R Studio (https://www.rstudio.com/) front-end and ggplot2 library(Wickham, 2009) to plot the graphs. Alpha values for statistical tests were chosen based on average sample size as follows: α = 0.05 1< N ≤ 10, α = 0.01 10 < N ≤ 100 and α = 0.001 100 < N

## Author contributions

IG and AE conceived the experiments and wrote the manuscript. VS carried out qRT-PCR analysis of mRNAs immunoprecipitated under stringent conditions. AK carried out the EmGFP-Tm1-I co-immunoprecipitation assays. The rest of the experiments and data analysis were carried out by IG.

## Acknowledgements

We thank Denise Montell for sharing unpublished data, antibodies and discussions. We thank Damian Brunner, Tze-Bin Chou, Elizabeth Gavis, Antoine Guichet, Daniel St Johnston and the Developmental Studies Hybridoma Bank for fly stocks and reagents, and the TRiP at Harvard Medical School (NIH/NIGMS R01-GM084947) for transgenic RNAi fly stocks used in this study. Stocks obtained from the Bloomington *Drosophila* Stock Center (NIH P40OD018537) were used in this study. We are grateful to Frank Wippich and Simon Bullock for his comments on the manuscript. We thank Sandra Müller and Alessandra Reversi for fly transgenesis, the EMBL Advanced Light Microscopy Facility and Leica for providing cutting-edge microscopy, and the EMBL Genomics Core Facility for their help with qRT-PCR. This work was funded by the EMBL.

---

**Table S1, related to Fig 1:**
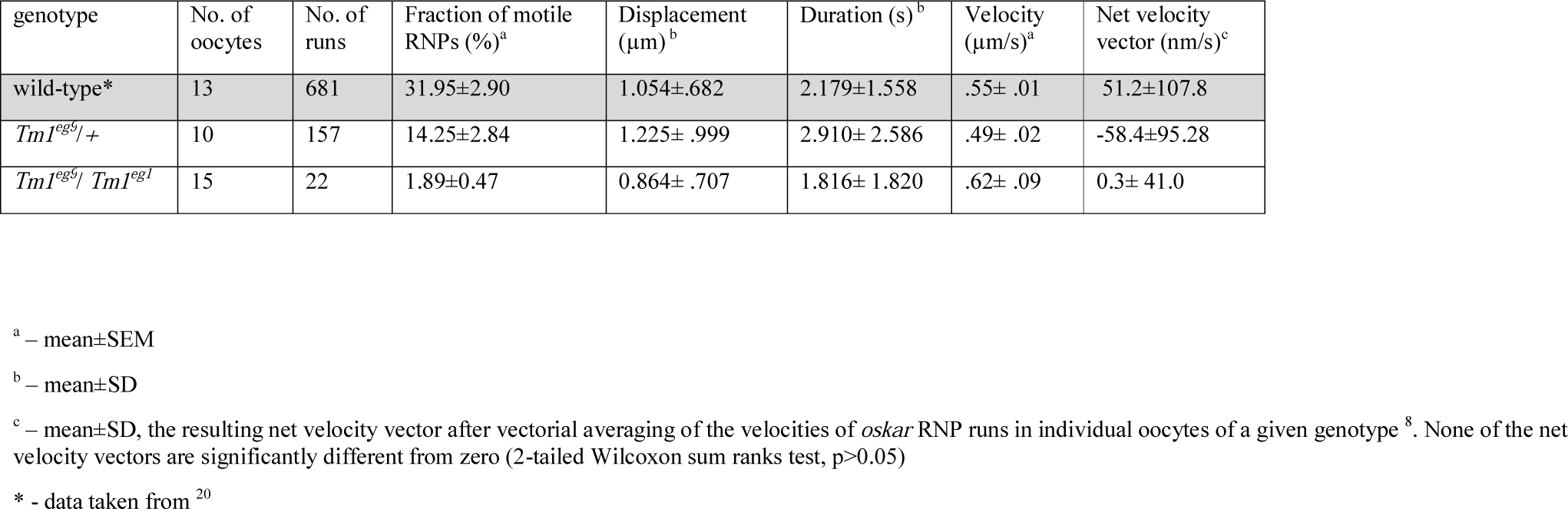
Motility statistics of *oskar* RNPs in *Tm1*^*gs*^ mutant oocytes *in vivo*

**Table S2:**
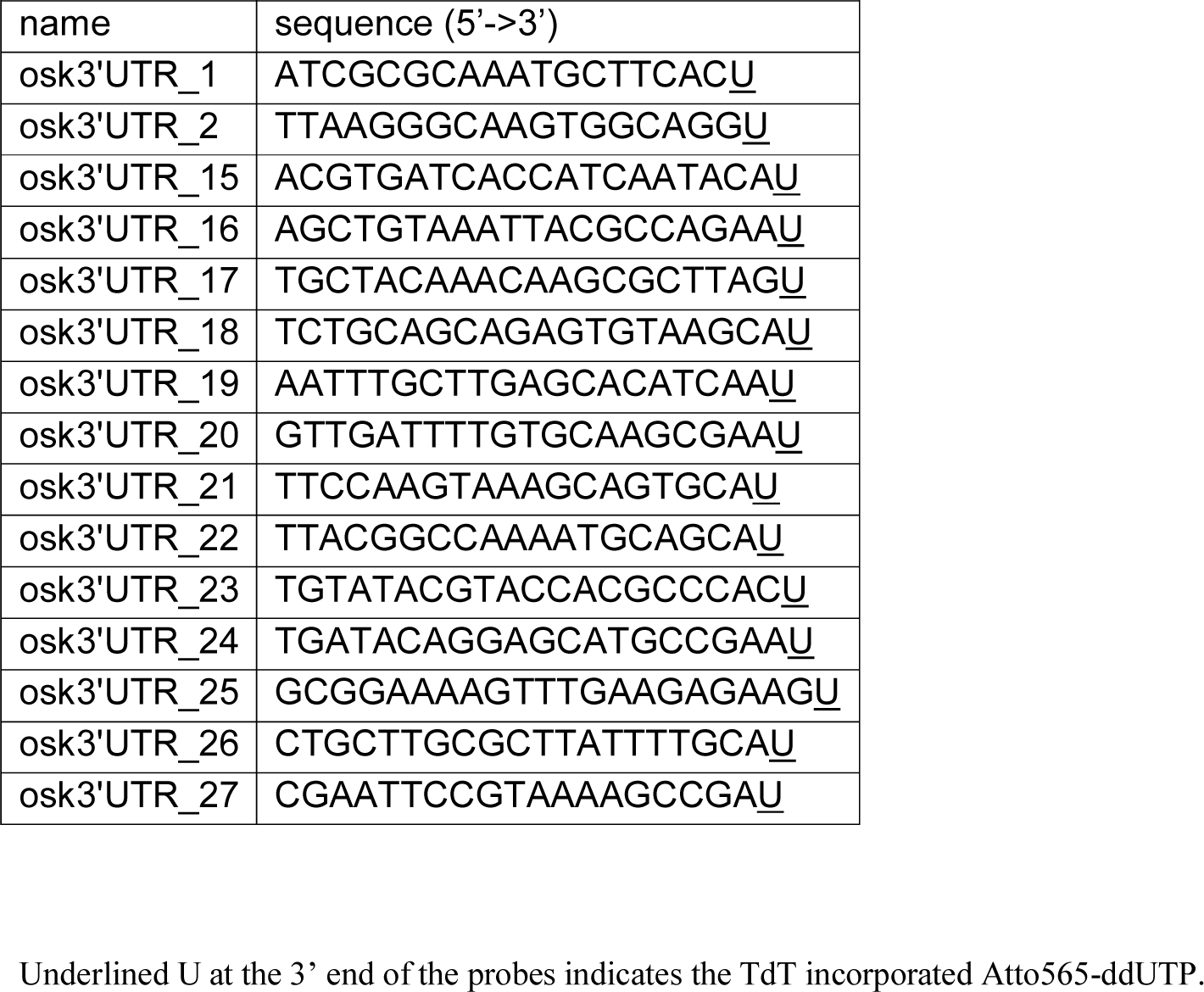
Sequence of smFISH probes targeting the oskar 3’ UTR.

